# Attention modulates the neural geometry of auditory representations

**DOI:** 10.1101/2025.10.09.681368

**Authors:** CA Pedersini, I Muukkonen, P Wikman

**Author notes:** Corresponding author: Caterina Annalaura Pedersini.

## Abstract

Selective attention enables the brain to prioritize behaviourally relevant information in complex environments, yet its neural mechanisms remain unclear. Here, we combined fMRI with representational geometry analyses to examine how attention reshapes neural population codes during naturalistic sound scenes. Participants attended to individual sound objects presented either with distractors from different categories (three objects across, 3OA) or the same category (three objects within, 3OW). Attention attracted neural population responses in low-dimensional manifolds toward the representation of each object presented in isolation, consistent with a reorganization of high-dimensional representational geometry. This effect depended on acoustic context: in 3OA, attended representations were significantly closer to object-alone representations than to distractors, whereas in 3OW attentional modulation was minimal. Category-specific effects were observed, with speech engaging left-lateralized language regions beyond the auditory cortex. Finally, stronger behavioural performance predicted greater representational attraction. These findings show that selective attention reshapes neural geometry in a context– and category-dependent manner closely linked to behaviour.

## Introduction

One of the fundamental feats of hearing is the ability to separate auditory scenes with multiple overlapping sound sources into their constituent auditory objects^1–6^. Selective attention significantly contributes to this process by boosting the neural processing of the relevant objects at the cost of the irrelevant sounds^7–9^. When the sound sources can be separated based on simple features, such as location, pitch or loudness, selective attention influences source separation in the auditory cortex (AC) by modulating the gain or tuning of responses in neurons processing the task relevant feature^10–19^. Yet, often in natural scenes, the objects cannot be separated based on simple features alone and, thus, it is thought that attention also operates using abstract object-level magnification^20–25^. The current literature on auditory object-level attention is dominated by studies of ‘cocktail-party’ speech, where multiple speakers are talking at the same time. Such studies show that neurons track attended speech more accurately than ignored speech^8,22,26,27^. While such models provide important insights into attentional modulation of speech, they do not fully capture the complexity of natural auditory environments, nor do they account for the dynamic mechanisms underlying attentional selection when this selection is defined by complex features of objects.

A promising strategy to address these limitations is to model neural activity within the framework of brain space manifolds^28^. These manifolds capture the low-dimensional geometry of distributed population responses^29^. Despite heterogeneous activity at the level of single neurons, empirical evidence suggests that population activity often lies on such manifolds, reflecting latent dynamics that constrain responses and shape behavior^30^. In this reduced brain space, the dominant axes of voxel covariation (population dimensions) become explicit, allowing us to directly characterize how cognitive states such as attention dynamically reshape the brain’s representational geometry^31–38^. Furthermore, this approach captures population-level processes that traditional decoding methods may overlook^39^. One central dynamic mechanism, which has been associated with selective attention, is *attraction*. An *attractor* is a point on the manifold which draws activity towards it, i.e. a valley in the representational landscape toward which nearby activity converges^31^. In the visual domain, selective attention has been shown to dynamically shift attractor points, warping representational manifolds toward goal-relevant objects^40,41^ and modulating semantic dimensions^42^ to enhance task-relevant categories. However, whether such dynamics generalize to human auditory attention in scenes with multiple natural objects remains elusive.

In our recent study^6^, we investigated selective attention in complex auditory scenes using fMRI. Participants listened to a diverse set (144 exemplars) of natural sound clips from three categories – speech, animals, and instruments – across three experiments (Fig. 1). In each experiment, participants focused on one of three simultaneously presented auditory objects. The overlapping objects were drawn either from different categories (*three objects across categories*, 3OA experiment) or from the same category (*three objects within category*, 3OW experiment). In addition, each object was presented alone (*object alone, OA* experiment). Using searchlight multivoxel pattern analysis (MVPA), we assessed whether local neural patterns elicited by complex auditory scenes resembled those elicited when each object was presented in isolation. We found that neural activation patterns of scenes more closely resembled those elicited by the attended objects than by either of the two distractors, indicating that selective attention modulates local representational structures of scenes^43,44^. Yet, while our results are compatible with dynamic brain space mechanisms discussed above (e.g., *attraction)*, the methodology we used could not reveal the precise geometric transformations underlying the attention effects. Consequently, in the current study, to disentangle these dynamics, we reanalyzed the neuroimaging data using a geometry-based framework of brain spaces.

**Fig. 1.**
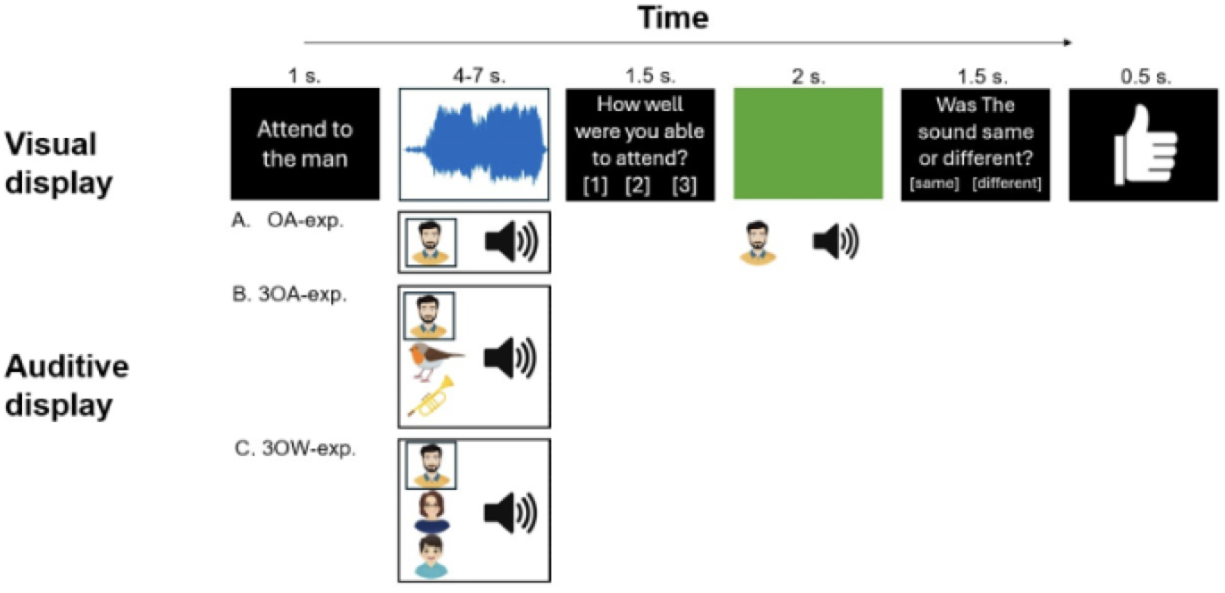
Behavioral paradigm in the three experiments (OA, 3OA, 3OW). Each trial begins with an instruction to attend to either an auditory display or a visual display, followed by stimulus presentation. The participant was presented with either one auditory object (OA), or three auditory objects (3OA or 3OW), as well as a visual video of a scrolling waveform display. Participants attended either to the designated sound object, while ignoring the video display (*attend sound object task*) or to the video display while ignoring the auditory stimulation (*visual control task*). Note that in 3OA and 3OW during the attend sound object task also the two distractor sounds were to be actively ignored. After each trial, participants rated their attentional success, and on some trials performed a match-to-sample task with feedback (see Wikman et al., 2025 for details).

Our pipeline implements two complementary alignment steps: first, we hyperalign (*Generalized Procrustes Analysis*^45–49^ all participants’ functional data into a shared information space (brain space); second, we use Procrustes alignment to map scene manifolds onto a reference space defined by *object-alone (OA)* representations. As attention may dynamically reshape scene manifolds either by altering the spacing of object representations or by rotating the dominant axes of population activity toward task-relevant features, we extracted two complementary geometric metrics: Euclidean Distance, capturing shifts in the spacing and clustering of object representations (*attraction*), and Principal Angle, assessing rotations of population axes toward task-relevant dimensions (*axis rotation*). These dynamics likely support complementary functions for downstream readout: *attraction* may refine object-level relationships, enabling more precise identification of the relevant object, while *axis rotations* may additionally provide coarse, category-level gain, particularly in cross-category scenes. While these mechanisms illustrate two complementary ways in which attention can reorganize representational geometry, they are not exhaustive; other transformations, such as changes in manifold curvature or dimensionality, may also contribute to the reorganization of population codes.

Building on this approach, we address three main questions: (1) how does selective auditory attention reshape the geometry of neural population spaces? (2) how does the surrounding sound scene modulate this geometry? and (3) how does the organization of neural geometry relate to behavior? We hypothesize that attention shifts the geometry of scene manifolds toward the object-alone brain space locations of the attended object of the scene compared to the distractor object representations. This effect is expected to depend on whether distractors belong to the same or a different category as the attended object^6^. We also expect that the organization of neural geometry in auditory regions will bear behavioral relevance, reflecting how attentional modulation of representational structure relates to subjective rating of attentional performance. By investigating these questions, we aim to reveal how top-down and scene-level factors interact to organize auditory scene representations at the population level.

## Results

We used three experiments to provide complementary insights into how attention shapes the geometry of brain spaces within naturalistic soundscapes. In each experiment we presented a diverse set (144 exemplars) of natural sound clips from three categories – speech (two male, two female and two child speakers), animals (cat, dog, monkey, songbird, seal, whale), and instruments (saxophone, trumpet, 12-string guitar, baritone guitar, jazz organ, classical organ). The *3OA* experiment tested how attention influences processing of sound objects when each object of the scene belongs to different categories (i.e., one speaker, one instrument and one animal object). The *3OW* experiment tested how attention affects processing of auditory scenes composed of objects from the same category. The *object-alone* experiment, in which objects were presented alone, served as a reference for two purposes: (1) aligning data across experiments using Procrustes analysis, and (2) computing distances between attention-modulated scene representations and object-alone representations (see below).

Our analytical approach is summarized in Figure 2. Briefly, for each participant, region of interest (ROI,^50^), and hemisphere we first constructed data matrices representing neural responses to individual sounds, either at object or exemplar level, with rows corresponding to individual objects (or exemplars depending on the analysis) and columns to vertices within each ROI. To align representational spaces across participants, we then applied Generalized Procrustes Analysis (GPA)^45,48^, which estimates orthogonal transformation matrices (rotations, translation and reflections) that minimize distances between functional patterns while preserving each participant’s intrinsic geometry, thereby aligning corresponding dimensions across participants (Fig. 2A). To reduce dimensionality and quantify the structure of condition-specific response patterns for each subject and ROI, we thereafter performed Principal Component Analysis (PCA) separately for each experiment on the aligned data. We extracted PCA scores and applied Procrustes alignment (without scaling) to map the representational spaces of experiments 3OA and 3OW (based on the attended and distractor objects of the scenes separately) onto the reference space of the OA experiment. This alignment ensured that the axes of the representational spaces were consistently oriented across experiments, so that distances and relationships between conditions reflected true neural differences rather than arbitrary rotations, reflections, or component orderings in the neural data (Fig. 2B, left panel). Finally, we quantified geometric changes using two complementary metrics: Euclidean distance (ED) and principal angle (PA). ED measured how strongly each scene representation in brain space was attracted toward the corresponding object-alone representation (i.e., the neural representation of each object when presented in isolation). PA quantified changes in manifold orientation, capturing rotations of the low-dimensional axes associated with attended versus distractor representations (Fig. 2B, right). To assess attentional effects on neural geometry, we compared ED and PA between attended and distractor coordinates/manifolds across all 360 cortical ROIs. To determine whether the observed differences exceeded chance levels, we applied the full analysis pipeline to surrogate data (see Supplementary Material).

**Fig. 2.**
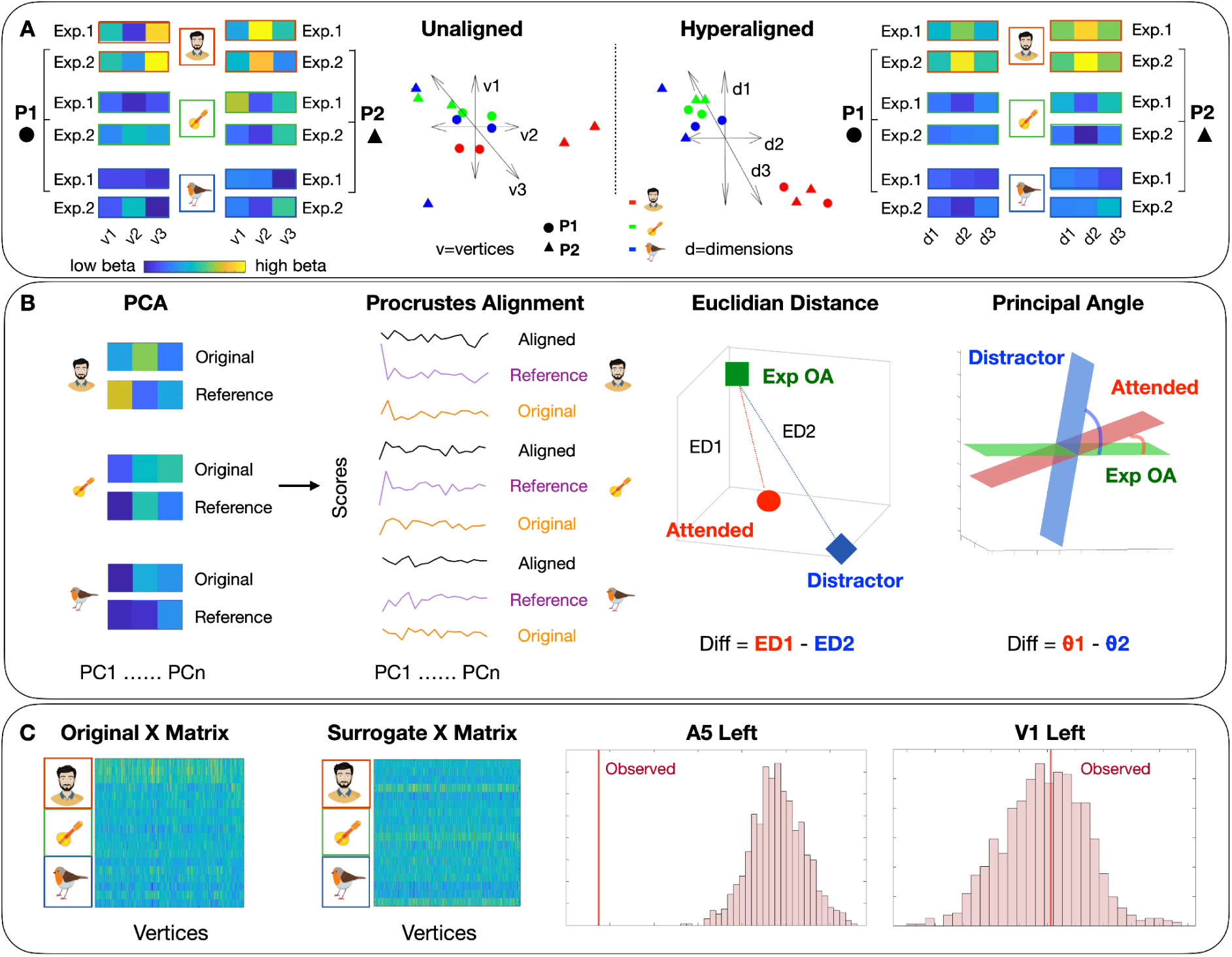
Overview of the geometry analysis pipeline. **A**. Hyperalignment of neural response matrices across participants using Generalized Procrustes Analysis. *Left:* fMRI Beta-estimate matrices from two participants for six stimuli (two per category) and their corresponding geometric representations in a three-vertex space. *Right:* Geometric representations in the aligned model space and the aligned beta-estimate matrices for the same participants and stimuli. Hyperalignment was performed to transform subject-specific responses from the anatomical space, defined along vertex axes (v1, v2, and v3), into a common, shared representational space across participants, defined along information dimensions (d1, d2, and d3). **B. Dimensionality reduction and comparison of representational geometry.** PCA was applied to the hyperaligned data across all experiments to reduce dimensionality. The resulting PC scores were then aligned to the scores from experiment OA (reference) using Procrustes analysis without scaling. This alignment is critical because the axes of the PCA space can differ across experiments arbitrarily; aligning the scores ensures that the geometry of the representational space is comparable, between the conditions, i.e. distances and relationships between conditions reflect true differences rather than arbitrary rotations or reflections of the axes. These aligned scores were then used to quantify attention-related changes in representational geometry by computing Euclidean distances (ED) and Principal Angles (PA). Specifically, ED and PA are computed for each condition of experiments 3OW and 3OA relative to experiment OA, and the differences between them are extracted. **C**. **Surrogate null distribution pipeline.** To estimate null distributions, stimuli of the same category across the three input matrices (attended 3OW, distractors 3OW, and experiment OA) were concatenated, their rows shuffled and then split back into separate matrices to generate structureless surrogate data. The same analysis pipeline as in B was applied to these surrogate matrices to compute null distributions of differences between attended and distractor stimuli. Observed differences were then compared to these null distributions to derive permutation-based p-values. These results can be found in the supplementary material.

We first conducted a whole-brain analysis across all 360 ROIs to localize the overall effects of attention on brain manifold geometry. Subsequent analyses focused on a subset of 34 ROIs per hemisphere, encompassing auditory, temporal, parietal, and frontal areas (see Fig. 6E for a spatial map of these regions). In the main text, results are reported using the first three PCs, explaining the highest variance. We also performed a variance-optimized analysis, selecting the number of PCs required to explain at least 70% of the variance in each dataset and using the maximum across all matrices to ensure consistent representation across experiments (see Figs. S3 and S5). Results from this analysis were largely consistent with the findings produced using 3 PCs.

### Attention reshapes the geometry of neural manifolds

We started with tackling two questions: (1) How does selective attention reshape the geometry of neural population spaces? and (2) How does the surrounding soundscape modulate this geometry? We quantified differences between attended and distractor conditions in terms of Euclidean distance and Principal angle, separately for experiments 3OA and 3OW.

Regarding the **Euclidean Distance**, we compared scene coordinates aligned to the attended object (*attended coordinates*) with those aligned to the distractor object (*distractor coordinates*) of the scenes. In experiment *3OA*, where the attended objects and distractor objects belonged to different categories, attention shifted manifold geometry toward the object-alone coordinate of the attended object of the scene. That is, *attended coordinates* were closer to those of the object alone coordinates than *distractor coordinates*. This effect was observed bilaterally in auditory and ventro-temporal regions for instruments and animals and extended to predominantly left frontal regions for speech (Fig. 3A). Surrogate analyses confirmed that these effects were significantly different from chance (Fig. S01). The visualization in Fig. 3B shows how the attended manifold gravitates toward the object-alone manifold, in comparison to distractor manifolds for two ROIs. In contrast, in experiment 3OW, where all scene objects belonged to the same sound category, no significant differences between *attended and distractor coordinates* were found (Fig. 3C). As can be seen in the visualization of category centroids in a 3D PCA space (PC1–PC3), both *attended and distractor coordinates* lie close to the object-alone configuration, with no systematic separation between the manifolds. Since in 3OW all objects of the scene were from the same category, it was fully expected that on a category-level there would not be any significant differences between the category centroids of the *attended manifolds* and the *distractor manifolds*. It is, however, important to note the distances were calculated at the object level (e.g., saxophone), not at the category level (e.g., instrument), and, thus, object-level changes may have been present in 3OW. Consequently, to assure that small differences in the category centroids between attended and distractor manifolds did not occlude more fine-grained within-category changes, we aligned the centroids of the OA, *attended*, and *distractor* brain space *manifolds* and recomputed the same distance metric (see *Methods – Geometric analysis of neural subspaces* for details, and Fig. 5A for a graphical illustration). This analysis, however, did not reveal significant attentional effects in 3OW.

**Figure 3.**
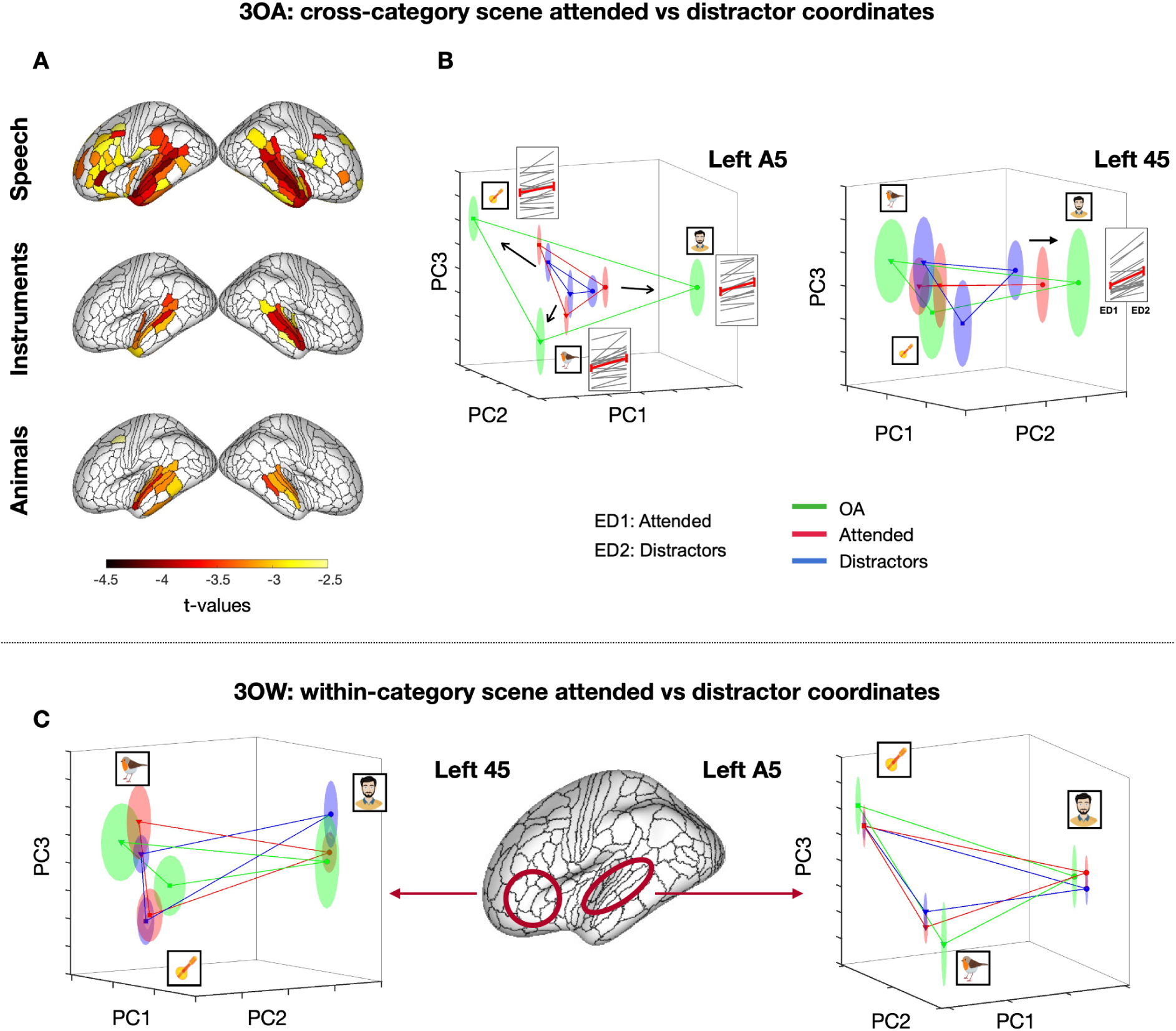
The effect of attention on the geometry of neural representations. A. The effect of attention on Euclidean distance in the 3OA experiment. Whole-brain t-values for speech, instrument, and animal sounds are displayed on lateral surfaces. Negative t-values indicate that scene representations when aligned based on the attended object of the scene are closer to the object-alone representational coordinates than scene representations when aligned based on the distractor of the scene. **B. Attentional effect of attraction.** Visualization of the mean centroids of three stimulus categories across experiments in a 3D PCA space (PC1–PC3) for two regions (Glasser Atlas) The black arrows in the Left A5 panel indicate the displacement of attended representations toward the object-alone representation. For each significant effect, Euclidean distances ED1 (attended vs. object alone) and ED2 (distractors vs. object alone) are plotted for individual participants (grey lines) and group averages (red line with SEM error bars) to illustrate the consistency of the effect. **C. Effect of attention on Euclidean distance in the 3OW experiment.** Visualization of the mean centroids of three stimulus categories across experiments in a 3D PCA space (PC1–PC3) for two ROIs Left 45 and A5. Ellipsoids around each centroid represent the standard error of the mean (SEM), and colored lines connect category centroids within each experiment (green: object alone; red: attended object, and blue: distractors). The lateral brain surface shows the location of the ROIs.

The fact that we found no changes for the 3OW experiment and clear changes for the 3OA experiment indicates that attention and sound-scene configuration together affect manifold dynamics –attentional attraction toward the object-alone configuration occurred only when attended and distractor objects belonged to different categories. As above, we thereafter tested whether the attraction effect in 3OA was dominated by category-level or more fine-grained within-category object level changes, and thus, we aligned all manifold centroids to that of OA. Yet, as for 3OW this analysis yielded no significant effects, indicating that the effect was driven by category-level changes (i.e., on the level of speech, animals, and instruments rather than on the level of the specific objects within the categories). Finally, we performed the same Euclidean Distance analyses as described above at the exemplar level (see *Construction and Alignment of Representational Spaces – Exemplar Level* in the Method section). However, these analyses did not reveal any significant effects, further indicating that the differences between attended and distractor manifolds in 3OA were dominated by category-level attraction.

Regarding the **Principal Angle**, in 3OA, a few left-hemisphere regions showed category-specific effects: FOP2 (frontal operculum) for speech and PBelt (posterior belt of the auditory cortex) for instruments, where the angle between object-alone manifolds and *attended manifolds* was smaller than that between object-alone manifolds and *distractor manifolds* (Fig. S2A). In 6v (ventral part of area 6) attention, in contrast, increased the angle for animal objects. FOP2 is implicated in higher-order auditory and speech-related processing^51^, PBelt in processing complex sounds such as music and environmental stimuli^52^, and 6v in sensorimotor integration and auditory-motor interactions^53^. On the contrary, no significant differences were observed in the 3OW experiment (Fig. S2B), consistent with the lack of attentional effects on Euclidean distance. These results suggest that, while attention consistently shifts neural representations toward the representations of the attended object of the scene when presented alone (OA), effects on the orientation of the neural manifold are more limited and region– or category-specific. Notably, including a higher number of PCs in the analysis did not substantially alter the Euclidean distance results, which remained similar with only a slight increase in the number of involved regions (Fig S3A). For Principal Angle, using more PCs revealed effects in different auditory (A1), parietal (LIPv), and frontal (6a and IFSp) regions, only for instruments and animals, confirming that the orientation of neural subspaces is region– and category-specific (Fig S3C).

### Distractor manifolds shift in relation to manifolds of ignored scenes

The shift of scene-related representations in brain space toward the attended object, relative to distractors, could arise through three non-mutually exclusive mechanisms: (i) attention may selectively enhance the neural representation of the attended object, (ii) attention may suppress the representations of distractor objects, or (iii) both processes may operate simultaneously. To distinguish between these possibilities, we conducted a post-hoc analysis to test whether *distractor coordinates* move closer or further away from the *object-alone coordinates* in comparison to coordinates estimated from scene presentations during the visual control task (see Fig 1, and methods: *Experimental session*), where all objects of the scene were fully ignored. This analysis was conducted for all ROIs that showed a significant difference in Euclidean distance for any of the sound categories in either hemisphere (FDR-corrected p < 0.05; Fig. 3B) in our main-analysis (see *Attention reshapes the geometry of neural manifolds*), including 67 ROIs per hemisphere.

This analysis revealed that *distractor coordinates* were consistently attracted toward their corresponding *object-alone coordinates* in both experiments, relative to when the same scenes were presented during the visual task that diverted attention from all sounds (Fig. 4). This effect was more extensive in 3OA (Fig. 4A), encompassing auditory, temporal, and frontal regions, whereas in 3OW (Fig. 4B) it was more restricted, involving primarily temporal and some frontal areas. Thus, it seems that attention attracts both *attended and distractor coordinates* towards their corresponding *object-alone coordinates,* albeit more strongly for attended than distractor manifolds (see *Attention reshapes the geometry of neural manifolds*). To further characterize these effects, we visualized category centroids in a 3D PCA space (PC1–PC3). In Experiment 3OA, manifolds during the visual control task were compressed toward the center of the representational space, showing a robust and widespread effect across regions. In contrast, in Experiment 3OW, visual control task manifolds were shifted closer to the object-alone centroid albeit further away from the OA-centroids compared to distractor centroids.

**Figure 4.**
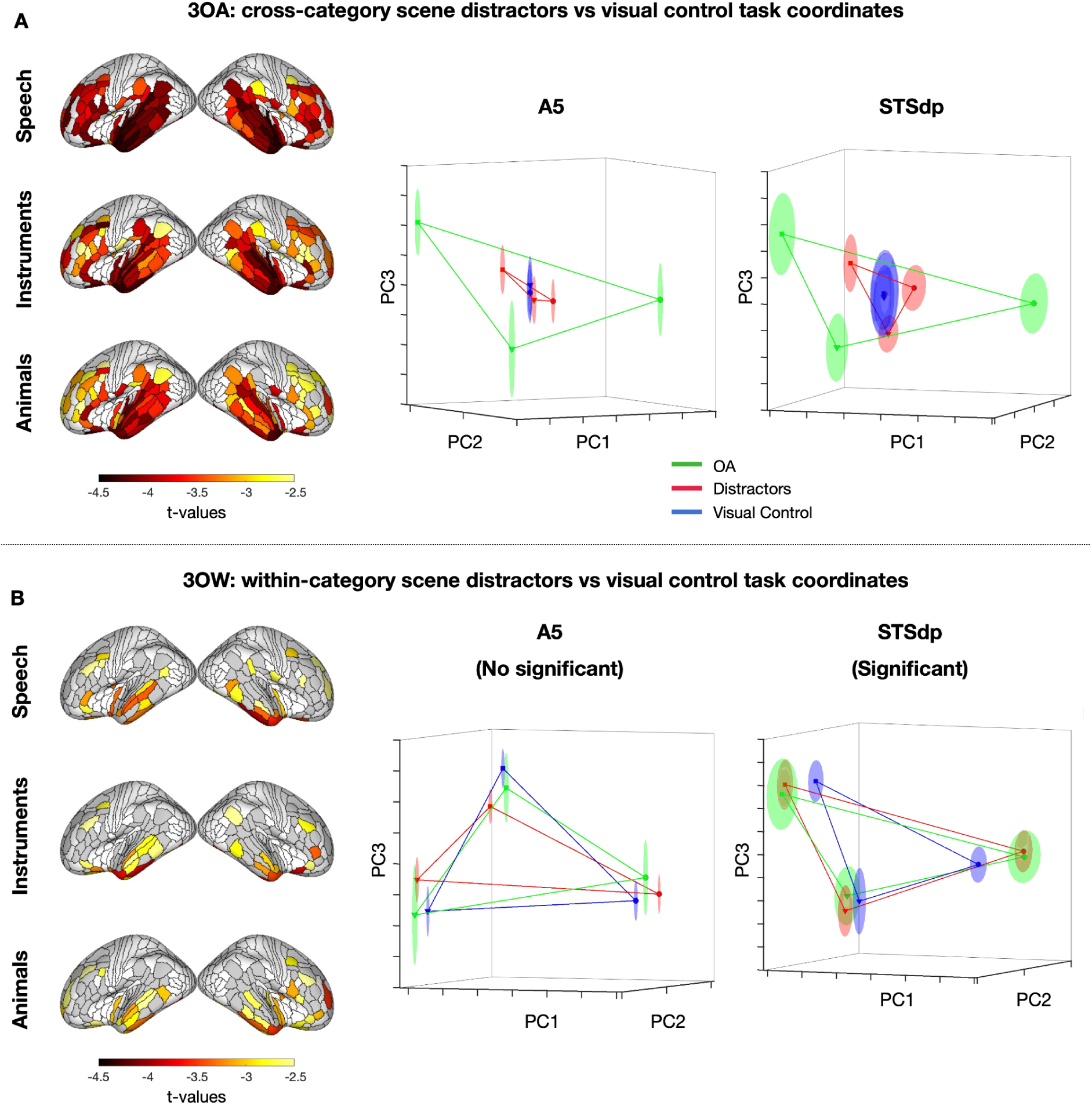
Shifts in Distractor Representations Relative to Ignored Scenes (visual control task), for experiment 3OA (A) and 3OW (B). Left: Effect of attention on Euclidean distance in the 3OA (top) and 3OW (bottom) experiments. Whole-brain *t*-values for speech, instrument, and animal sounds are displayed on lateral surfaces. Negative *t*-values indicate that distractor manifolds are closer to the object-alone coordinates than similar scenes during the visual control task (all sounds ignored). Gray-colored ROIs indicate the spatial locations of all brain regions included in the analysis. **Right: Attentional effect of attraction.** Mean centroids of the three stimulus categories are visualized in a 3D PCA space (PC1–PC3) for two example regions: left A5 (high-order auditory region) and STSdp, (the dorsal-posterior subdivision of the superior temporal sulcus). Ellipsoids around each centroid represent the standard error of the mean (SEM), and colored lines connect category centroids within each experiment (green: object alone; red: distractors; blue: visual control task).

### Neural Geometry of scenes changes according to subjective evaluation of task performance

After each trial participants gave a rating of their ability to selectively attend to the specified object of the scene. This behavioral data were extensively analyzed in our previous work (Wikman et al., 2025), where we reported that the ratings showed significant effects for both Experiment (OA, 3OW, 3OA) and Category (speech, instruments, animals), with the lowest ratings observed in experiment 3OW and for instrument sounds (for similar analyses on the match to sample task, see also Wikman et al., 2025). Building on these findings, we wanted to evaluate whether participants’ perception of their ability to attend to the target object affects neural representational geometry.

We found that in 3OW subjective ratings significantly predicted Euclidean distances, primarily in the superior temporal sulcus (STS). As participants reported a greater ability to attend to the target of the scene, the distance between attended and object-alone representational points decreased, with nearly all participants showing negative slopes. Notably, this relationship emerged only after centering the PC scores by subtracting the category-specific centroid, thereby aligning each stimulus representation to its *object-alone (OA)* centroid and removing general category-level differences (see Fig. 5A for a schematic of the centroid alignment and Fig. 5B for the corresponding spatial map of significant effects). This procedure allowed us to isolate the gravitational effect independent of broader category-level structure. At an uncorrected threshold (p < 0.01), the effect extended beyond the left STS to include a few other areas including area A5, indicating a subtle, yet consistent influence of attentional performance on auditory scene representations (Fig. S4A). In experiment 3OA, uncorrected results (p < 0.01) likewise revealed an effect in the right A1 (Fig. S4B). Despite this effect being very consistent across participants, the effect did not survive FDR correction, likely due to the presence of many regions showing weak or no effects. Importantly, including a larger number of PCs in the analysis did not substantially change the results, and significant FDR-corrected effects were only observed for experiment 3OW after centroid alignment (Fig. S5).

**Fig. 5.**
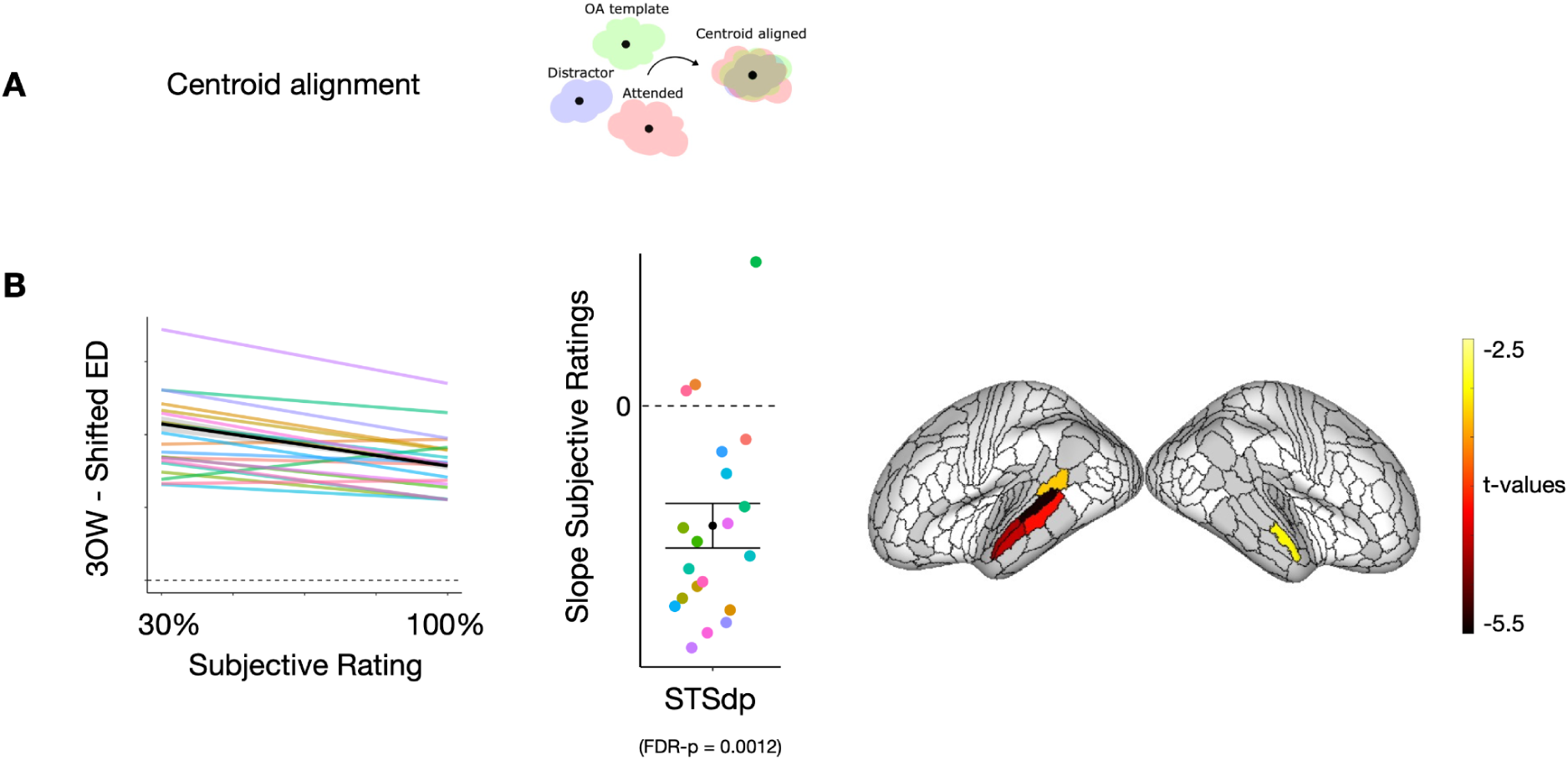
Subjective rating predicts Euclidean Distance (ED). A. Graphical representation of the centroid alignment procedure. **B**. **Regions showing significant effect of subjective rating on Euclidean Distance.** Spatial map of brain regions where subjective ratings of ability to attend to the target object of the scene significantly predicted ED, showing a general trend across all scenes–i.e., the distance between the attended and the object-alone representations decreased as subjective rating increased. We show the results for 3OW, after centering the PC scores by subtracting the category-specific centroid, thereby aligning scores to the OA centroid. Gray-colored ROIs indicate the spatial location of all brain regions included in the analysis. For a significant ROI (left STSdp) we display: (1) slopes for individual participants (colored lines) together with the group mean (black line) ± 95% Confidence Interval, and (2) the mean slope value ± Standard Error with individual subject values as colored dots.

### Comparison of Acoustic and Neural Manifolds

The *object-alone* experiment offers the opportunity to trace the geometry of auditory representations along the auditory ventral pathway. Here, we specifically asked whether neural manifolds evolve from acoustically grounded geometries in early auditory regions towards more compact and clustered configurations in higher-level auditory cortex and semantic networks. To address this, we quantified the alignment between acoustic and neural manifolds using the Procrustes Distance as a measure of alignment between spaces^33,54^. A systematic pattern emerged in the distribution of Procrustes Distances, with smaller distances to acoustic features (pitch, spectral centroid, harmonic-to-noise ratio, amplitude modulation standard deviation and frequency content standard deviation, see *Stimulus acoustic features*) in primary and secondary auditory regions compared to temporal, parietal, and frontal areas (fig. 6E). This indicates that neural representational geometries converge more closely toward the acoustic feature manifold than would be expected by chance only in primary and secondary auditory regions (fig. 6C). To illustrate these effects, we visualized the data projected into a 3D PCA space (PC1–PC3) for a few representative regions (Fig. 6D): MBelt, A4, and STSva as examples of regions showing a similar pattern between acoustic and neural manifolds (lower Procrustes Distance), and TPOJ2, 45 and TE2p as examples of regions exhibiting more divergent patterns (higher Procrustes Distance). These regions were chosen to exemplify the two main effects observed: (1) TE2p (inferior temporal cortex) showed extensive overlap among stimulus representations, with both acoustic and categorical distinctions largely lost; and (2) TPOJ2 and 45, representing parietal and frontal areas, showed a categorical structure, primarily separating speech from the other categories.

**Fig. 6.**
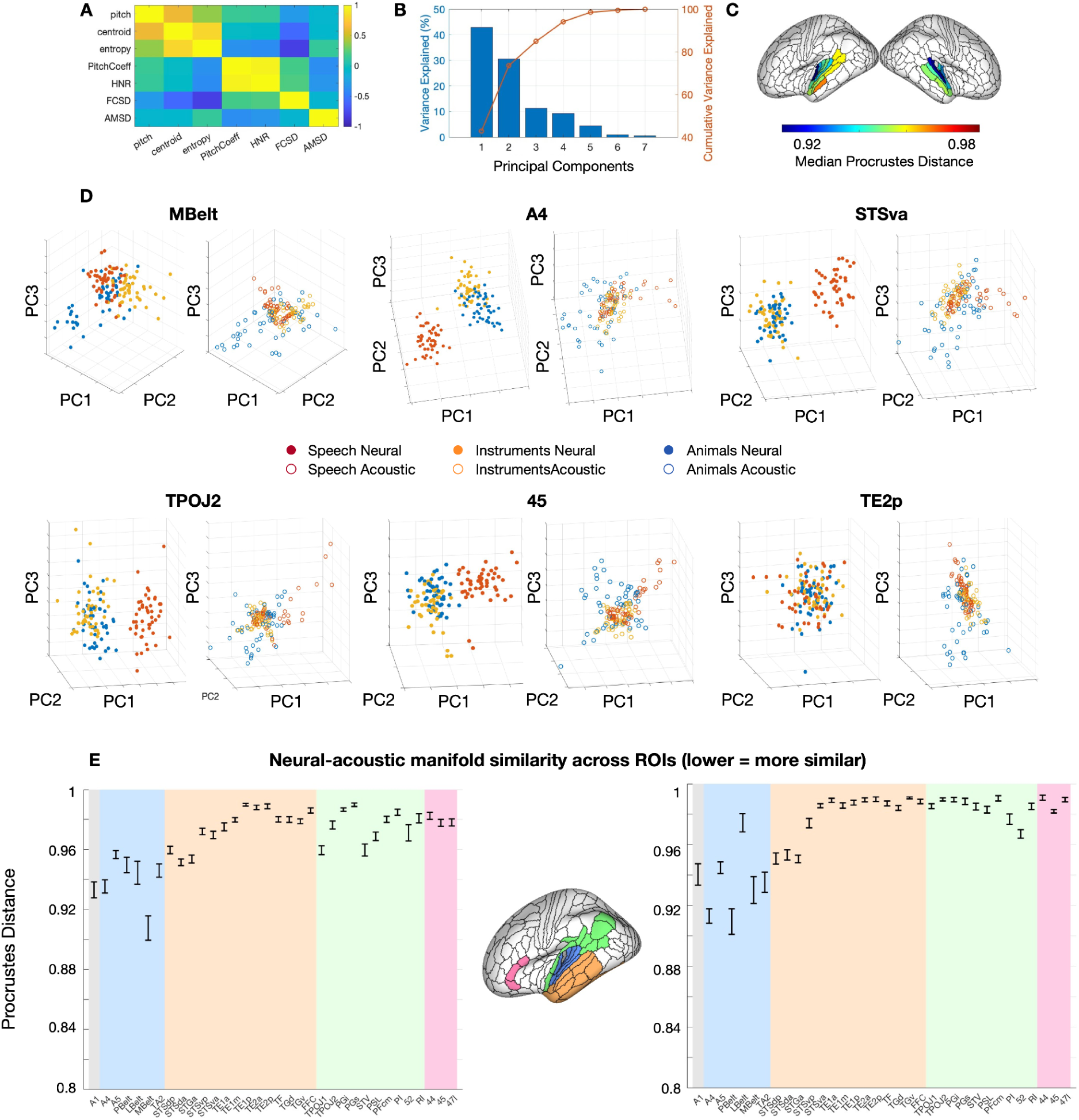
Similarity between acoustic and neural manifolds. **A**. Correlation matrix between acoustic features of the 144 auditory stimuli. **B**. Variance Explained and Cumulative Variance Explained across Principal Components. **C.** Spatial map showing the distribution of median Procrustes Distance between the acoustic and neural manifold, significantly different from chance (p<0.05 – 10000 permutations). **D**. Projection of the 144 stimuli onto the first three principal components, color-coded by category, for the neural manifold of experiment OA and the aligned acoustic manifold (Procrustes alignment with scaling), shown as the median across participants. Represented regions are selected from the left hemisphere as examples of low (MBelt, A4, STSva) and high (TPOJ2, 45, TE2p) Procrustes Distances. **E.** Neural–acoustic manifold similarity (median ± SEM) across 34 regions in the left and right hemispheres. Regions are grouped into five clusters (also shown on the left lateral brain surface): primary auditory cortex A1 (grey), secondary auditory regions (blue), temporal regions (orange), parietal regions (green), and frontal regions (pink).

## Discussion

In the present study, we used human fMRI data to examine how selective attention shapes neural population geometry of auditory scenes. Our main findings can be summarized into four key points. First, brain space geometry analyses showed that attention reshapes scene manifolds – shifting scene representations toward the representational points of the scene constituting objects (estimated from trials where each object was presented in isolation, object-alone). Second, this effect was stronger for the attended object than the distractor objects of the scene, but only when the objects constituting the scene were from different sound categories (3OA). Third, stronger subjective attentional performance was associated with closer alignment between attended and object-alone representations, particularly in auditory and ventro-temporal regions. Fourth, comparing acoustic and neural manifolds revealed a systematic representational progression: primary and secondary auditory regions closely reflected acoustic structure, whereas higher-level temporal and frontal regions showed increasingly divergent and abstract geometries. Together, these findings highlight the dynamic interplay between sensory structure, attention, and behavior in shaping auditory population geometry.

### Attention attracts scene representations towards attended-object representations context-dependently

The current study was inspired by our previous results demonstrating that scene representations are dominated by fMRI activation patterns of the attended object of the scene^6^. Based on these results we conjectured that two viable population dynamics could be at play: *attraction* or *axis rotation*. The results of the current study showed clear evidence for attraction and only marginal evidence for axis rotation, and consequently our discussion will focus on attraction. In brain space, attraction can be conceptualized as certain points of the neural manifold behaving as attractors, i.e., valleys in the landscape towards which nearby activity evolves^28,29^. Previous vision studies have shown selective attention to act by dynamically shifting attractor points, causing manifolds to collapse towards goal-relevant objects in brain space^41,42^. Our representational geometry analyses provide support for similar mechanisms in human audition, i.e., attention reshaped the population space of overlapping scenes by gravitating/attracting neural responses of scenes toward the representational points of the sounds when presented alone. This attention-induced ‘geometric attraction’ complements our previous MVPA results^6^: while MVPA quantifies increased discriminability along readout dimensions, geometry analyses reveal how attention reorganizes the entire representational landscape, drawing population activity toward the attended object of the scene. Thus, we argue that attraction is a central neural mechanism by which attention ‘increases the contrast’ of task relevant objects in scenes. Attraction as a key attentional mechanism is also a viable candidate to classic ‘gain-increase’, where attention increases the gain of neurons encoding the attended stimulus features^55,56^. Notably, unlike gain-increase, attraction can account for how attention modulates processing of natural objects, represented by neuronal patterns embedded in parallelly active populations of neurons.

In classic attractor models, however, population dynamics converge near to a finite set of stable fixed points on the manifold, with activity outside these attractors transiently decaying toward one of them^28^. Yet, scene representations gravitated toward the object-alone patterns but remained distinct, rather than collapsing onto the same brain-space points. We still argue that this pattern is compatible with an attraction model of attention due to the following reasons: (1) Attention may fluctuate in a pendulum-like fashion between the target object and distractors over the presentation of the scene and consequently, the fMRI pattern reflects a time-averaged mixture of neural states with a net bias toward the attended object (akin to alternation dynamics observed in binocular rivalry^41^). Consistent with this notion, our participants reported that their attention was not continuously fixed on the target sound of the scene^6^. (2) Attentional attractor points are probably not the exact object-alone states. Instead, they are probably latent points in brain space that adjust dynamically to behavioral goals and to the ongoing estimate of the target’s identity (see *Attraction depends on subjective attentional performance*). (3) Instead of specific points, attractors may be a continuous set of equally stable states, i.e., a manifold of fixed points^57–60^. Thus, the attention induced attractors may be lines oriented toward the attended object, or a ring encircling the object-alone representation. In this view, attention may bias positions along the manifold or tilt the landscape, rather than forcing convergence to a single discrete state.

Crucially, we observed that the **surrounding soundscape** modulated attraction in brain spaces: in 3OA scenes, where the attended and distractor sounds belonged to different sound categories (speech, instruments and animals), attention selectively attracted scene representations toward the object-alone representation of the attended object of the scene, relative to distractors. Importantly, this effect reflected categorical differences: when we aligned the centroids of attended and distractor manifolds to the object-alone template, no effects remained significant. This may indicate that when the scene contains sounds from distinct categories, attention reshapes neural geometry primarily relative to the category-level structure, rather than on the level of individual sound objects or exemplars. Alternatively, the ‘Category’ vector may have emerged as dominant, overshadowing finer within-category shifts, since the geometric distance between categories is much larger than the distance between instances within a category. This does not, however, indicate that individual objects or exemplars cannot act as attractor points for cross-category scenes. Rather, as described in the hierarchical-decomposition model of attention^61^, the *dominant* level of attraction may change hierarchically over time, that is, cross-category scenes may first gravitate to the category-level manifold, then the object-level and finally to a specific exemplar-level. Thus, if we would have presented our scenes for longer than 4-7 seconds, our results may have differed.

In contrast to cross category scenes, within-category (3OW) scene representations classified according to the attended or distractor object of the scene were equally close to the category-level object-alone manifold. This was indeed expected, as in such scenes, attraction cannot occur at the categorical level, since all stimuli belong to the same category. Instead, we expected attention to gravitate specific scene representations towards the attended sub-categorical points of individual objects. Yet, in 3OW we did not observe scene representations to move closer to the attended objects than the distractor objects. This lack of results may indicate that geometric distances between objects within the same category (e.g., Violin vs. Guitar) are inherently much smaller than between categories making the attentional shift too subtle to detect in the main group-average contrast due to fMRI noise levels. However, the result is also consistent with previous notions that object-level representations may exist in such sparse populations of neurons that they cannot be estimated using fMRI^52^. Alternatively, the opposite is also possible, namely that object-level representations transverse several macro-anatomical fields along the auditory cortex, thus eluding our ROI-based approach^62^. Finally, it is possible that object-level attraction did not emerge in the current study because object-specific manifold structure needs to be gradually learned (see^63–65^ or reinforced over time (see *Attraction depends on subjective attentional performance*). Taken together, our findings suggest that selective attention reshapes neural population geometry in a category– and context-dependent manner.

### Distractor representations shift relative to fully ignored scenes

Selective attention has been found to not only modulate the processing of task relevant sounds but also actively ignored (distracting) sounds^66,67^. Many models posit that the processing of distractors is actively suppressed, thus, complementing the enhanced processing of attended sounds^67–70^. This pattern of results has been observed for simple sound stimuli, yet direct evidence for neural suppression of *natural complex distractor objects* is sparse. In the current study, we observed the opposite of suppression for distractor objects, i.e., the representation of the distractors in the scenes was enhanced rather than suppressed. That is, distractor manifolds were closer to single object templates than manifolds of similar scenes with all objects ignored (participants focused on a visual task and ignored the auditory scenes altogether). Thus, our results indicate that the state pattern of distractors were enhanced in scene representations compared to fully ignored scenes. Furthermore, this pattern was observed for both within category scenes (3OW) and cross-category scenes (3OA). These results may emerge in two ways: (1) attention may enhance the representation of all objects of the scene, albeit more strongly for the target object; (2) as discussed above, participants could not always fully focus their attention on the target, causing fluctuation between target–distractor states in brain space (see *Attention attracts scene representations towards attended objects context-dependently*). Thus, our data argue against a pure distractor-suppression account and suggest instead that the neural processing of distractors is likely context and goal dependent^8,66^.

### Attraction depends on subjective attentional performance

There is longstanding interest in linking neural changes to behavioral performance. In the context of neural manifolds, this has motivated efforts to examine whether variations in geometric properties – such as distances or alignment within neural representational spaces – systematically relate to behavior^31,35,71^. Building on this framework, we tested whether the observed attraction of scene representations toward the attended object of the scene changed based on subjective behavioral performance indicators. Specifically, we tested whether subjective ratings of attentional performance (the ability to focus attention on the target sound and ignore the distractors) predicted the Euclidean distance between neural manifolds.

Our results highlight attentional performance as a key factor shaping neural representational geometry: higher subjective ratings of attentional performance predicted smaller Euclidean distances between attended and object-alone *exemplar* representations. Notably, in within-category scenes (3OW) behavioral effects emerged only after centroid alignment, indicating that attentional engagement predicts fine-grained, within-category shifts in the neural manifold – subtle changes in how individual exemplars are represented relative to their category centroid. In cross category scenes (3OA), a similar effect was only observed in the primary auditory cortex (A1) and was evident in both raw distances and after centroid alignment (note that this effect did not survive FDR-correction). This suggests that, even in cross-category scenes, behavioral performance affected simple sound features rather than overall category-level distances (e.g., frequency representations, see^72^. It is also possible that the category-level attraction in *3OA* emerged very rapidly, leaving little room for behavioral effects to enhance it. It is important to note that participants had not heard any of the exemplars used before the scene tasks. Thus, the behavioral attraction results are consistent with our conclusion that the auditory cortex continuously updates an internal model of the heard sound as the task unfolds, allowing exemplar-level representations to emerge as attractor points for scenes (see above). Taken together our findings demonstrate that the ‘geometric attraction’ of population responses indeed has behavioral relevance by reflecting participants’ subjective estimates of their level of attentional performance.

### Sound category dependent attraction emerges in both acoustically grounded and higher-level ventral stream regions

Beyond scene-context-dependent effects (see, *Attention attracts scene representations towards attended objects context-dependently*) attention also operated differently for the three sound categories (speech, instruments and animals). For speech, attentional modulation was observed not only in auditory and temporal regions (core, belt, Superior Temporal Sulcus – STS) but also in extensive left-lateralized frontal areas, associated with higher-order language processing and speech comprehension^73,74^. In contrast, for instruments and animals, attentional effects were more restricted to auditory and temporal regions (core, belt, STS), reflecting that the focal point of attentional modulation being of spectral and intermediate-level auditory representations^75,76^. This is consistent with views arguing that non-speech objects are mainly encoded in the same subfields that encode features that differentiate them from each other^77^. Yet, we want to highlight that in 3OA attentional attraction was also observed in middle temporal regions for both instruments and animals, the temporal pole for instruments and lateral inferior temporal cortex for animals. Thus, attraction effects in these regions may indicate that attention also operated to some degree on supramodal or semantic representations^78,79^ of animals and instruments.

The functional specialization of different fields of the auditory cortex and auditory ventral stream regions beyond the auditory cortex has been debated^75,80^. Therefore, we also evaluated the functional role of different regions showing attentional attraction in a data-driven manner: by comparing acoustic (reflecting different spectral and temporal features of sounds) and neural manifolds, we directly addressed how auditory representations transform along the ventral stream. Our findings reveal a systematic transformation of representational geometry, from early acoustic encoding to more abstract, higher-order representations along the ventral stream. We observed lower Procrustes distances between acoustic and neural manifolds in primary and secondary auditory regions, as well as in the superior temporal sulcus (STS), indicating neural manifolds that partially preserve the geometry of the acoustic feature space^81,82^. Notably, in the acoustic manifold, animal sounds were more widely dispersed along the principal components, whereas speech and instrument sounds occupy more overlapping regions. This broader acoustic spread progressively diminished along the cortical hierarchy, indicating that these fine-grained acoustic distinctions are lost as neural representations become more abstract.

Higher-order temporal, parietal, and frontal regions exhibited progressively larger Procrustes distances, reflecting a reorganization of neural manifolds away from the low-dimensional acoustic representation. Crucially, in our results, higher Procrustes distances do not necessarily indicate stronger categorical coding; rather, they reflect greater divergence from the acoustic geometry. Categorical organization emerges in secondary auditory regions (A4), where all three categories of sounds occupy almost fully distinct regions of the space (see Fig 6). Moving beyond A4, to STS the Procrustes distances remain relatively low and stimuli show a clearer separation mainly between speech and other sounds (instruments and animals). Parietal and frontal areas show partial categorical structure, while ventro-temporal regions (inferior temporal cortex) exhibit extensive overlap among stimulus representations, with both acoustic and categorical distinctions largely lost. These results align with previous findings that anterior and posterior superior temporal regions differentiate speech, music, and other sound categories^52,75,77,83^, highlighting a progression from feature-based to more abstract population codes along the auditory hierarchy.

A similar hierarchical transformation has been observed in the visual system along the ventral pathway for object recognition. Early sensory regions encode detailed stimulus features in high-dimensional population activity, whereas downstream areas reduce dimensionality and cluster responses into more abstract, robust representations^84,85^. This framework combines the efficient coding hypothesis^86^ and the observation that higher-order neural codes occupy low-dimensional subspaces, supporting reliable computations despite noise^87^. High-dimensional population codes also exhibit a characteristic power-law structure, ensuring smooth and robust encoding of stimulus details^88^. These findings suggest that hierarchical representational transformations are a general principle of cortical sensory processing. Our geometric approach extends this framework to audition, providing a quantitative characterization of how auditory representations evolve along the ventral pathway, from acoustically grounded population codes to more abstract manifolds. This transformation occurs in both early and higher-order auditory areas, although it remains unclear which specific aspects of the representation are modulated by attention.

Taken together, the comparison between acoustic and brain-space manifolds suggests that attentional attraction engages both regions whose geometry primarily reflects physical acoustic features and regions whose geometry is decoupled from physical properties. Importantly, this does not by itself show that attention altered the predominant information content in those spaces: the same region may co-represent acoustic and abstract information, with the dominant representation shifting over time and with context. Thus, determining which specific information dimensions attention modulates in each subfield will require targeted, time-resolved analyses in future work.

### Future directions

The geometry of brain spaces can be altered through multiple mechanisms. In the present study, the predominant effect we observed was that selective attention induced gravitation/attraction of neural population responses to auditory scenes toward the object-alone representation. We, however, also examined whether the manifolds of scenes classified by the attended object of the scene was more aligned with object-alone manifolds than the manifolds classified based on the distractors of the scene (axis rotation), which we quantified as the angle between the main axes of the manifolds. Surprisingly, only a small number of regions showed significant effects. Notably, our analyses focused exclusively on between-space differences—that is, the geometric relationship between attended, distractor, and object-alone manifolds—without investigating within-space geometry, such as the angle between objects of the same category. Previous work suggests that selective attention may reshape the internal structure of neural manifolds by collapsing or expanding representations of individual objects within a category^89^. For example, the internal structure of the attended manifold may become more like that of the object-alone configuration than to distractor manifolds. Future work could examine these within-space effects to provide a more comprehensive account of how attention reshapes representational geometry.

A second consideration concerns spatiotemporal resolution: our results lack a temporal dimension. Yet, attention exhibits key temporal dynamics—initial separation of target sounds from distractors^6^ followed by selective gating of semantic/linguistic processing for exclusively the attended input^43^. Temporal dimensions are also an important component in the hierarchical-decomposition model^61^. Consequently, future work with high-temporal-resolution recordings such as EEG/MEG could reveal both within-space effects and the temporal unfolding of attentional modulation across hierarchical sound processing stages. Furthermore, given fMRI’s limited spatial resolution, future intracranial data are required to fully characterize attention-induced attraction in brain spaces. Nonetheless, our current results offer a strong starting point, by indicating that attraction effects are widespread along the ventral processing stream, and they depend on both the scene type and the sound category of the attended object.

### Conclusions

Our study demonstrates that selective attention reshapes neural population codes in a context– and category-dependent manner, effectively ‘attracting’ representations of a soundscape towards the representation of the attended sound object. This effect was modulated by the surrounding soundscape, with stronger attentional modulation when distractors belong to different categories (3OA), and varies across stimulus types, engaging left frontal regions for speech but remaining largely confined to auditory areas for non-speech sounds. Importantly, these geometric changes are behaviorally meaningful: participants’ subjective attention ratings predict the proximity of attended representations to their object-alone templates. Together, these findings highlight the functional relevance of representational geometry in auditory perception and provide a population-level framework for understanding how attention prioritizes behaviorally relevant information in complex environments. Future work could explore how these mechanisms interact with other cognitive processes such as learning and memory, and whether similar geometric principles govern attentional selection in other sensory modalities or real-world listening contexts.

## Method

### Participants

Twenty healthy right-handed monolingually native Finnish speaking participants underwent an fMRI session (8 females, age range 19–32, mean age 22.7 years, standard deviation 3.68). All participants were students at the University of Helsinki or Aalto University, had self-reported normal hearing, normal or corrected-to-normal vision and no history of developmental or neurological disorders. We assessed handedness by using the Edinburgh Handedness Inventory^90^ and acquired written consent from the participants before the study.

### Ethics statement

Participants were compensated monetarily €15/h for their participation in the study. Written informed consent was obtained for the sharing of processed anonymized data from each participant. The experiment was accepted by the University of Helsinki Ethical Review Board in the Humanities and Social and Behavioral Sciences, and we conducted the study in accordance with the Declaration of Helsinki.

### Stimuli

The experiment included auditory and visual stimuli, presented with Presentation 24.0 software (Neurobehavioral systems, Berkeley, California, USA). The auditory stimuli were presented binaurally through earphones (Sensimetrics model S14; Sensimetrics, Malden, Massachusetts, USA) and auditory volume was adjusted individually for each participant at the beginning of the session at ca. 85 dB. Visual stimuli were projected onto a mirror attached to the head coil. The participant answered the questions with an fMRI-compatible response pad with either the index-, middle– or ring finger of their right hand.

Auditory stimuli comprised sound clips from three object categories: speech, animal and instrument sounds, each with six subcategories. The speech subcategories included vocalizations from two adult females, two adult males, and two children (one girl and one boy). Animal sounds were sampled from huskies, cats, whales, monkeys, birds, and seals, while instrument sounds came from a baritone guitar, a 12-string guitar, a saxophone, a trumpet, an organ, and a bebop organ. Each subcategory contained eight exemplars, resulting in 144 distinct auditory clips (3 categories × 6 subcategories × 8 exemplars). Sound durations ranged from 4 to 7 seconds, with an average of 5.3 seconds. Speech samples were taken from emotionally neutral spoken dialogues recorded in previous studies^91,92^ and segmented into 4–7 s clips, each containing one sentence (Audacity 3.3.3). Animal sounds were sourced from free online databases, selected for variability in acoustic features (e.g., a howling husky rather than a monotonous bark) and minimal background noise. Recordings were denoised in Logic Pro X using the Waves X-Noise plugin (20–50 dB reduction). Instrument sounds were generated from the same speakers as the speech samples, pairing voice types with instruments of similar frequency ranges (e.g., low male voices with baritone guitar). Dialogue pitch and duration were analyzed in Logic Pro X (Melodyne plugin), adjusted to the 12-Tone Equal Temperament scale, and converted to MIDI. Outlier frequencies were corrected and resulting audio files were volume– and timbre-matched through compression in Logic Pro X and Audacity (threshold = −30 dB, ratio = 5:1, attack = 1.11 s, release = 1.0 s). Instrument samples were drawn from Logic Pro X and Spectrasonics Omnisphere, with organ sounds from the Logic Vintage B3 Organ plugin. All auditory stimuli were prepared using the same procedure as in our previous study^93^, and further details on preprocessing, loudness normalization, and filtering can be found therein.

Visual stimuli were generated following the procedure described in detail in^93^. Briefly, scrolling waveform videos were created from all auditory clips (including training and non-target items; *N* = 214) using MATLAB (R2020a), with a one-second blue waveform window on a white background. Videos exceeded the duration of the corresponding audio by one second of silence and were compiled at 25 fps (MPEG4 codec, 2000 kbps bitrate) using *ffmpeg*. After screening for redundancy, 180 visually distinct control videos were selected for presentation.

### Stimulus acoustic features

For each exemplar sound, six acoustic features were extracted using MIRtoolbox (v1.8.1;^94^) in MATLAB (R2023b), yielding one value per feature and clip. Pitch was computed with *mirpitch* using the default autocorrelation method, selecting the dominant periodic component within 75–10,000 Hz. The spectral centroid (*mircentroid*) was defined as the mean frequency of the stimulus spectrum. The harmonic-to-noise ratio (HNR) was calculated as the ratio of the dominant periodic component (maximum autocorrelation) to the aperiodic signal component (https://www.fon.hum.uva.nl/praat/)^52^. The amplitude modulation standard deviation (AMSD) quantified temporal amplitude variability and was computed as the standard deviation of the rectified waveform after resampling to 60 Hz. The frequency content standard deviation (FCSD) captures spectral variability as the standard deviation of power across frequencies. Finally, spectral entropy (*mirentropy*) was computed as the relative Shannon entropy of the stimulus spectrum.

### Experimental session

Each participant completed three experiments within the same session. All the experiments followed an identical trial structure consisting of four phases (Fig. 1). First, an instruction was presented for 1 second, indicating whether the participant should perform an auditory or visual task. In auditory trials, the instruction also specified the sound object to attend to (e.g., “attend to the man”). This was followed by the stimulus presentation phase, during which the designated auditory or visual object was presented. In experiments with overlapping sound, the attended sound object was always presented 250-500 ms before the distractors. Simultaneously, a visual video clip (randomly sampled without replacement) was shown for the same duration as the auditory stimulus (4–7 seconds). After the stimulus presentation, participants provided a subjective rating indicating how well they were able to attend to the designated object. Responses were given within a 1.5 second window by pressing a button to select one of three options: 1 (less than 33% of the time), 2 (33–66% of the time), or 3 (more than 66% of the time). In 25% of the auditory attention trials, a match-to-sample task followed, indicated by a green visual display background. In this task, participants heard a 2-second sound sample and indicated whether it matched the attended sound object. The sample was either an exemplar of the attended sound object (target trial, 50% probability) or a different exemplar of the same object category that had not been used as the attended stimulus (nontarget trial). In 100% of the visual attention trials, the participant was presented with a short sample (2 sec.) of either the video clip they had been attending to (target trial, 50% probability) or another video clip (nontarget trial). The intertrial interval was randomized to last from 201 to 5500 msec and the response time window was 1.5 seconds. After answering, the participant received feedback on their performance (0.5 sec.; correct/incorrect).

In the **Object alone (OA) experiment,** participants were instructed to attend to the single auditory object presented in each trial. Although visual videos were displayed concurrently, they were never task-relevant, and participants were instructed to ignore them. Each run consisted of 36 trials, including two repetitions of the same sound object within the run, using different exemplars. The experiment comprised four runs in total, allowing for the presentation of all 144 sound object exemplars once, 48 for each category (speech, animals, instruments). Trial order was randomized for each participant.

In the **3OA experiment**, each auditory scene consisted of three sound objects belonging to different categories (e.g., boy, husky, and trumpet). Within each run, the participant performed two tasks: the *attend sound object task*, including 75% of the trials, and *the visual control task*, including the remaining 25% of the trials. In the former, participants had to attend to a pre-specified sound object, while the other two objects acted as distractors and had to be actively blocked out. The combinations of sound stimuli were randomized for each participant and systematically rotated to ensure balanced pairings (see Wikman et al., 2025 for further details on the pairings). Each sound object type appeared twice as attended across two different scenes, using a different exemplar each time. This resulted in 36 auditory attention trials per run (12 scene types × 3 attention conditions), with 108 unique exemplars presented once per run. In the latter, participants were instructed to fully ignore all auditory stimulation and attend to the visual video. As in the auditory task, the same 12 scene types were used for the visual control task, but with different exemplars, bringing the total to 48 trials per run (around 8 minutes). While the scenes remained the same across runs for the same participant, their order was randomized.

In the **3OW experiment**, each auditory scene consisted of three sound objects belonging to the same category (e.g., Male1, Boy, Female2 or Bird, Husky, Whale). As in the 3OA experiment, each sound object type was presented in two different auditory scenes. For each participant, the sound object types within a category were first randomly shuffled, then grouped so that each sound object appeared in exactly two scenes. This resulted in four unique scene subsets per category (speech, animals and instruments). Each sound object could serve in three roles—attended, distractor 1, or distractor 2—across the four scene types and three categories, yielding 36 auditory attention trials per run (3 attended objects × 4 sound scene types × 3 sound object categories). As in the 3OA experiment, the same 12 scene types were also used for the visual control task, but with different exemplars. See figure 1 (bottom section)^93^, providing more details on experiments and stimuli used.

Each participant completed a training session outside the scanner on the same day as the fMRI acquisition, during which each experiment was practiced for a total of two runs (including exemplars not used in the actual fMRI experiment). The fMRI session always began with the *object-alone* experiment, while the order of the remaining tasks was counterbalanced across participants, resulting in a total of 12 runs per subject. Short breaks were provided between runs, during which participants could rest and communicate with the researchers if needed.

### fMRI data acquisition

For a detailed description of the fMRI acquisition^93^. We report the parameters used in brief in Table 1.

**Table 1.** MRI-acquisition parameters used in the f/MRI data collection.

|  | TR | TE | Flip Angle | Voxel Matrix | Volumes | Slice Thickness | FOV | Slices | Resolution |
| --- | --- | --- | --- | --- | --- | --- | --- | --- | --- |
| **EPI** | 1.05 s | 30 ms | 75° | 78 x 78 | 360 (OA)<br>570 (3OA/3OW) | 2.7 mm | 21 cm | 45 | 2.7 mm × 2.7 mm × 2.7 mm |
| **T1** | 2.53 s | 3.3 ms | - | 256 × 256 | - | 1 mm | - | - | 1 mm × 1 mm × 1 mm |

### Preprocessing and analysis of fMRI data

Functional MRI data were preprocessed using FSL Version 6.00, part of FSL (FMRIB’s Software Library, http://www.fmrib.ox.ac.uk/fsl), including using MCFLIRT (Jenkinson et al., 2002), interleaved slice-timing correction, non-brain removal with BET^95^, and high-pass temporal filtering (with a cut-off of 100 Hz). Functional images were registered to the individual structural (T1-weighted) images using FLIRT^96,97^. All further fMRI analyses were conducted in surface space. Functional data were projected to the FreeSurfer average cortical surface (fsaverage7) using *mri_vol2surf*^98^, and all analyses were performed separately for each hemisphere on the surface.

For the first-level analysis, we used the preprocessed functional images as input and included nuisance regressors to account for motion and physiological noise. Specifically, we included the six standard motion parameters (three translations: *trans_x, trans_y, trans_z*; and three rotations: *rot_x, rot_y, rot_z*), framewise displacement (FD), and the first five components from the anatomical CompCor method (aCompCor) derived from signals in white matter and cerebrospinal fluid. These confounds were extracted from the output of fMRIPrep 20.2.5^99^ and entered as additional nuisance regressors in the general linear model using FSL FEAT.

In the first-level analyses, we fitted a general linear model (GLM) to the time series data of each vertex in each run, using FEAT at the subject level. All GLMs included separate boxcar regressors, defined with onset at the beginning of the first sound and duration covering the entire auditory stimulation (all three sounds), convolved with the canonical hemodynamic response function (HRF). For each of the three sound categories (speech, instrument, and animal), six distinct objects were used, and each was repeated twice within a run. This resulted in a total of 36 regressors (3 categories × 6 objects × 2 exemplars) in experiment OA, and 48 regressors in experiments 3OW and 3OA, corresponding to 36 auditory and 12 visual regressors. Nuisance regressors were included to account for all timepoints associated with the match-to-sample task. All further fMRI analyses were conducted at the level of individual ROIs, using the Glasser Atlas^50^.

### Construction and Alignment of Representational Spaces

To assess attention-driven differences in brain space geometry, we extracted one input matrix per experiment and condition for each ROI. The primary analysis was carried out at the object level, with supplementary analyses at the exemplar level to evaluate fine-grained alterations in scene manifold geometry.

#### Object level

For each subject, COPEs (contrasts of parameter estimates), representing the estimated effect size at each vertex, were first averaged across repetitions of the same sound object within each run and across runs. As described in the *Experimental Session*, in experiments 3OW and 3OA, each sound object served as a target in two soundscapes and as a distractor in four. To balance the attended scene matrix and distractor scene matrix, two COPEs were randomly selected without replacement from the distractor matrix for each object in each run. For each subject, ROI, and hemisphere, this yielded five data matrices, composed of 18 rows (one per unique sound object; six objects per category across three categories) and a number of columns equal to the number of vertices in the ROI. The five matrices corresponded to 3OW attended, 3OA attended, OA, 3OW distractor, and 3OA distractor conditions. In addition, we extracted two visual-condition matrices (one per experiment) representing the sound scenes when they were ignored. These matrices also had 18 rows, one per unique sound object, consistent with the auditory stimuli.

These matrices were first mean-centered by subtracting the mean activation across conditions for each vertex and then hyperaligned across participants using Generalized Procrustes Analysis (GPA)^45,48^ without scaling (see figure 2A for a graphical representation). This procedure returns transformation matrices that optimally align each subject’s ROI-level representational space to a shared, high-dimensional common space. GPA derives orthogonal transformation matrices (rigid rotations, translation and reflections) that minimize distances between functional patterns while preserving the representational geometry of each participants’ data. This procedure functionally aligns corresponding dimensions across participants. To reduce the dimensionality of the representational brain space and to quantify the structure of condition-specific response patterns for each subject and ROI, we applied Principal Component Analysis (PCA; *pca.m* function in matlab), on the aligned data from each experiment separately (see figure 2B for a graphical representation). We extracted the PCA scores and computed the cumulative explained variance across components. Once the PCA scores were extracted, we applied Procrustes alignment without scaling to map the representational spaces of experiments 3OW and 3OA (attended and distractor conditions) onto the reference space of experiment OA. This alignment ensures that the axes of the representational spaces are oriented consistently across experiments, so that distances and relationships between conditions reflect true differences in neural activity rather than arbitrary rotations, reflections, or ordering of the principal components. Scaling was omitted in both alignment steps to preserve the intrinsic geometry of the neural representational space, ensuring that distances between conditions reflect meaningful patterns of neural activity.

#### Exemplar level

At the exemplar level, individual exemplars were treated as separate conditions. Experiment OA included 144 trials (1 repetition per exemplar), while 3OA and 3OW each included 36 unique auditory trials of sound triplets in each run, either from different categories (3OA) or the same category (3OW), plus 12 visual-control trials. Importantly, the 3OA and 3OW contain only a subset of the total 144 exemplars, pseudorandomly selected and different across participants (see^6^ for details. For each subject, ROI, and hemisphere, this yielded five data matrices: 3OW attended (36 rows), 3OA attended (36), object-alone (144), 3OW distractors (72; 36 per distractor), and 3OA distractors (72; 36 per distractor), plus two visual-condition matrices (36 rows each). Because exemplars differed across participants in 3OW and 3OA, GPA was applied only to the OA matrices to derive a reference space. PCA was then performed per experiment, and the PCA scores of corresponding exemplars in 3OW and 3OA were aligned to the OA reference space using Procrustes alignment without scaling, to ensure a common alignment for cross-experiment comparisons.

### Geometric analysis of neural subspaces

To quantify differences between conditions in the geometry of the low-dimensional representation for each category, we extracted two metrics from the data projected to the low-dimensional PCs: Euclidean Distance (ED) and Principal Angle (PA).

#### Euclidean Distance

Euclidean distance was calculated between pairs of the same object (or exemplar) by comparing the PCA scores of experiment OA (reference) with the aligned scores of attended and distractors from 3OW or 3OA. Distances were then averaged across objects (or exemplars) within each category to yield a mean distance representing the overall category-level separation (see Fig. 2 for a graphical representation of the pipeline). To assess whether observed differences reflected within-category representational changes in addition to global shifts in category position, we performed a centroid alignment in some analyses (see Fig. 5A for a schematic description): for each subject and category, neural patterns from 3OW or 3OA were aligned to OA by calculating the vector connecting the category centroids and shifting all object-level representations to the same centroid while preserving the relative geometry within each category. EDs were computed using either centroid-aligned or unaligned data.

#### Principal Angle

Principal Angle was extracted to quantify how attention affects the orientation and the alignment of neural manifolds^32,34,100^. Specifically, we adapted the pipeline described by Panichello et al. (2021), to calculate the principal angle between category-specific subspaces, at subject-level. In brief, we applied a second PCA separately for each stimulus category and experiment (input matrix = 6 objects x 3PCs). This yielded eigenvectors (basis functions) of size 3×3. The first two eigenvectors (2×2) were used to define the plane-of-best-fit to the points defined by the rows of the 6 objects belonging to the same category and they explained more than 93% of the variance of each set of points in the 3D space. Given two vectors (*v1,v2*) defining the first plane and (*v3,v4*) defining the second plane, the cosine of the angle between the planes was computed from the determinant of the dot-product matrix between the spanning vectors, normalized by the determinants within each subspace (see^32^ for details). This yielded a measure ranging from 0° (identical subspaces) to 90° (orthogonal subspaces).

#### Statistical Analysis

To quantify attentional shifts on the geometry of brain representations, we compared ED and PA between scenes aligned based on the attended object of the scene and scenes aligned based on the distractor objects of the scene, including all ROIs (180 for each hemisphere) using a non-parametric sign-flipping permutation test. For each category, we computed the observed difference in representational dissimilarity (e.g., experiment 3OW attended vs. distractors), then generated a null distribution by randomly flipping the difference signs across 50,000 permutations. A permutation-based t-statistic was obtained as the ratio of the observed mean difference to the standard deviation of the null distribution, and two-tailed p-values were calculated as the proportion of permuted values exceeding the observed difference. P-values were extracted for ROIs (180 in each hemisphere) and metric (ED and PA) and corrected for multiple comparisons using FDR. We plotted significant results on a medial and lateral brain surface, overlapped with the Glasser Atlas.

To test whether the observed differences in representational geometry exceeded what would be expected by chance, we also applied our full analysis pipeline to surrogate data (see figure S1 for a graphical representation). We generated these surrogates by concatenating stimuli of the same category across the three input matrices (i.e., attended 3OW, distractors 3OW and experiment OA), shuffling the rows, and then splitting them back into separate matrices. This procedure preserves the overall covariance structure within each category while disrupting the specific arrangement of stimuli in the low-dimensional manifold. Applying the pipeline to the shuffled matrices produced null distributions of differences between attended and distractor conditions over 1,000 iterations, for each ROI and hemisphere. We then compared the observed differences to these null distributions to compute permutation-based p-values and z-scores. Finally, we extracted p-values at the whole-brain level, applied false discovery rate (FDR) correction across ROIs and hemispheres, and visualized the significant effects on the cortical surface, highlighting regions with reliably altered representational geometry.

### Relating Neural Geometry according to subjective evaluation of task performance

The goal of this analysis was to determine whether the effect of attention on the geometry of brain representations could be predicted by participants’ self-reported ability to focus on the designated object (Subjective Rating). In this case, Euclidean distances were extracted at exemplar level (see *Construction and Alignment of Representational Spaces: Exemplar level*), separately for each run. Distances were computed using either the raw values or after centroid alignment, as explained above. These trial-level distances served as dependent variables in our linear models. Data were filtered by condition (attended), and analyses were performed separately for each ROI (34) and hemisphere. For each subject, we estimated the slope of the relationship between subjective ratings and Euclidean distance using a linear regression model, implemented in RStudio (version 2024.12.1+563) running R (version 4.5.0; R Core Team, 2025), with the *lm* function from the stats package. The model included Subjective Rating (predictor of interest), Category (predictor of no interest), and their interaction term:

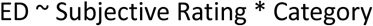

These slopes were then tested against zero using one-sample, one-tailed *t*-tests (*stats:t.test*), reflecting the a-priori hypothesis of a negative relationship between attention and representational distance (higher rating corresponding to lower distance). Resulting *p*-values were corrected for multiple comparisons across ROIs and hemispheres using the false discovery rate (FDR). To visualize the results, we plotted individual subject slopes as well as model-based mean regression lines with 95% confidence intervals.

### Comparison of Acoustic and Neural Manifolds

To examine the spatial distribution of similarity between the neural and the acoustic feature manifold, we implemented the following analysis pipeline. First, we derived an acoustic manifold by performing PCA on the six standardized acoustic features extracted from each exemplar sound. This yielded principal component scores representing the acoustic space. We then aligned the acoustic scores (3 PCs) to the neural scores (hyperaligned across participants) obtained from the object-alone experiment at exemplar level (144 exemplars), using Procrustes alignment with scaling. In this case, scaling was applied to account for differences in units and overall magnitude between the two spaces. From the aligned matrices, we extracted the Procrustes Distance – which represents the sum of squared differences between two sets of points after optimal scaling, rotation, and translation – between the acoustic and the neural manifold, separately for each ROI (34) and hemisphere^33,54,101^. Finally, we created a null distribution of distances and tested in which brain regions the neural representation geometry is closer to the acoustic feature manifold than would be expected by chance (p<0.05, 10000 permutations). To visualize the spatial distribution of the significant results, we plotted the significant Procrustes Distances on a lateral brain surface.

## Acknowledgment

This work was supported by The Research Council of Finland (grant number #1348353). The funders had no role in study design, data collection and analysis, decision to publish, or preparation of the manuscript. We want to thank M. Salmikivi, O. Varis, W. Vikatmaa, S. Rossow, and L. Lehtimäki for helping with data acquisition, as well as Ville Laaksonen and Jaakko Kauramäki for help with the study design.

## Data Availability Statement

All preprocessed behavioral and fMRI data, anonymized to protect participant privacy, as well as the corresponding analysis code will be deposited on the Open Science Framework (https://doi.org/10.17605/OSF.IO/AGXTH).

## Notes

https://doi.org/10.17605/OSF.IO/AGXTH)

### Competing Interest Statement

The authors have declared no competing interest.

### Summary of Updates

The primary updates are in the Discussion. While the main analysis pipeline is unchanged from the previous version, we have improved the clarity of the methods and provided a more in-depth discussion of the results. Moreover, we added additional results to more clearly highlight the key findings of the study.

