## Supplementary Material for "Attention modulates the neural geometry of auditory representations"


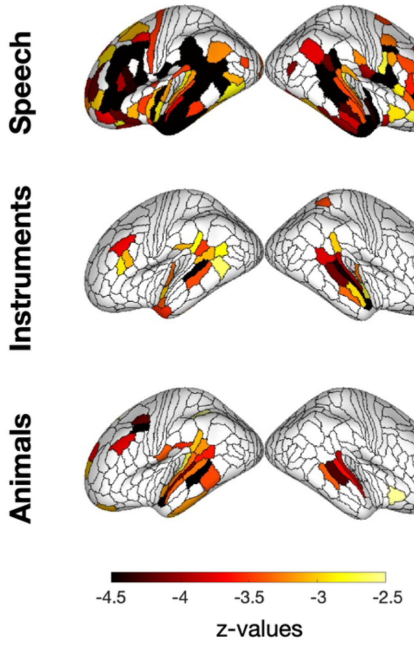


**Figure S1. Surrogate Analysis - Statistical significance relative to chance.** Whole-brain z-values show regions where the attentional effect on Euclidean distance differs significantly from a surrogate null distribution. Larger negative values indicate that the observed difference is significantly more negative than expected by chance. Importantly, regions showing significant differences in representational geometry (t-values see Fig 3) largely overlap with the significant z-scores obtained from the surrogate maps, indicating that these effects are robust and unlikely to arise from random stimulus-level structure.

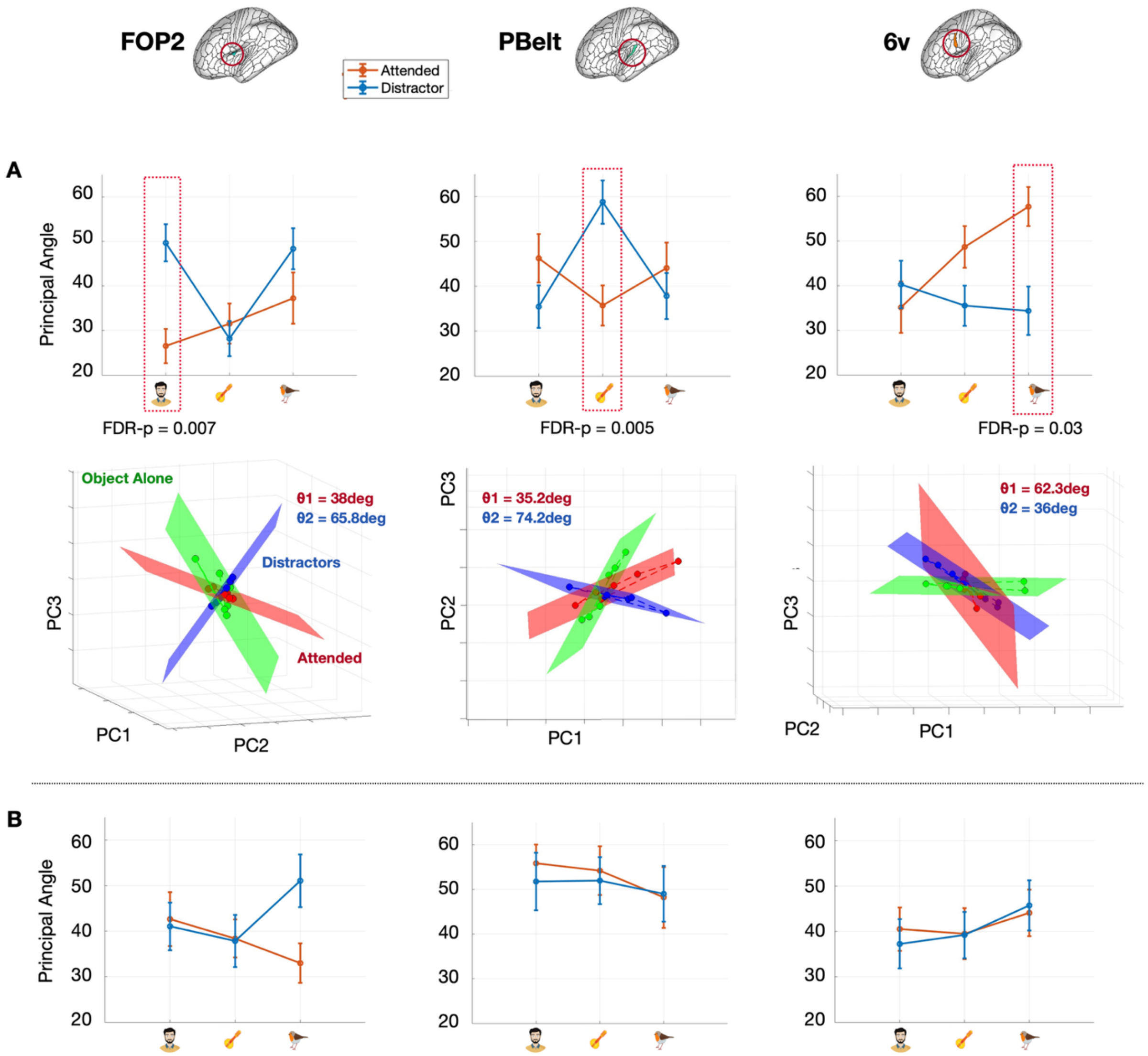


**Figure S2. Effect of attention on the geometry of neural representations. A. Effect of attention on Principal Angle in the 3OA experiment.** Upper panel: Principal angle as a function of category (speech, instruments, and animals) for attended (red) and distractor (blue) conditions, in ROIs showing an FDR-corrected significant attentional effect relative to chance. Bottom panel: Visualization of the three main planes extracted from the object-alone (green), attended (red), and distractor (blue) conditions in a 3D PCA space (PC1–PC3) for a representative subject. $\Theta_1$ indicates the angle between attended and object-alone, while $\Theta_2$ indicates the angle between distractor and object-alone. Dots represent stimulus exemplars projected onto each condition's 2D subspace, and dashed lines connect exemplars of the same category. **B. Effect of attention on Principal Angle in the 3OW experiment.** There were no FDR corrected significant effects in 3OW. We show principal angle as a function of category (speech, instruments, and animals) for attended (red) and distractor (blue) conditions for the regions showing significant effects in 3OA. The lateral brain surface in A indicates the spatial location of each ROI.

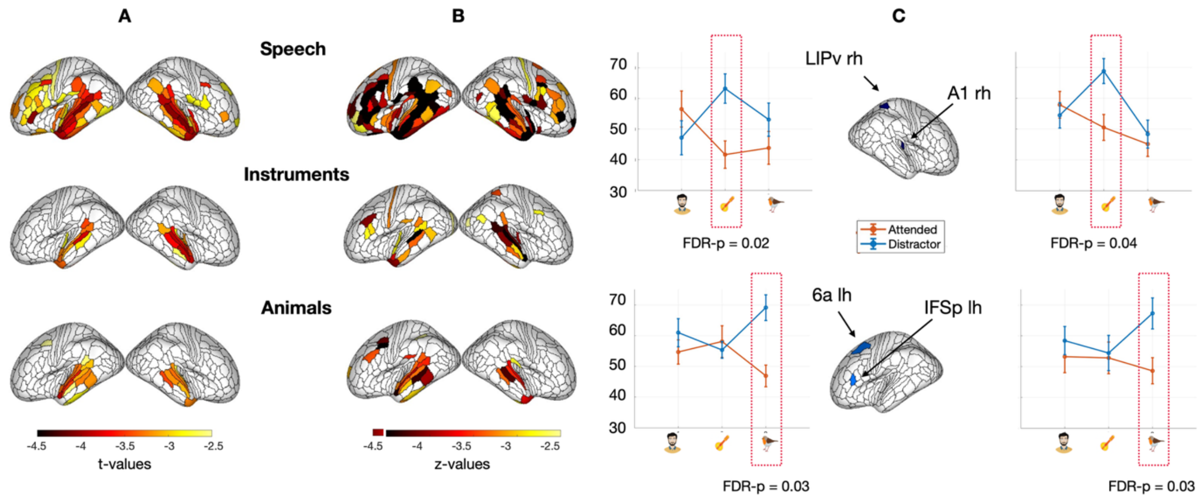


**Figure S3. Effect of attention on the geometry of neural representations in the 3OA experiment, with a variance explained of at least 70%. A. Effect of attention on Euclidean distance.** Whole-brain FDR corrected significant empirical results (t-values) displayed on the lateral surface for speech, instrument, and animal sounds. Negative t-values indicate that attended sounds are represented closer to the object alone than distractor sounds. **B.** Whole-brain results (z-values) showing regions in which the attentional effect on Euclidean Distance significantly differs from chance. Higher negative values indicate stronger deviations from chance, with the observed difference significantly more negative than the null distribution. Note that in both A and B the results are in line with the results computed on 3 PCs (Fig X and Y). **C. Effect of attention on Principal Angle.** Principal angle as a function of category (speech, instruments and animals) for attended (red) and distractors (blue), in the ROIs showing an effect of attention significantly different from chance. The spatial location of each region is shown on a lateral brain surface.

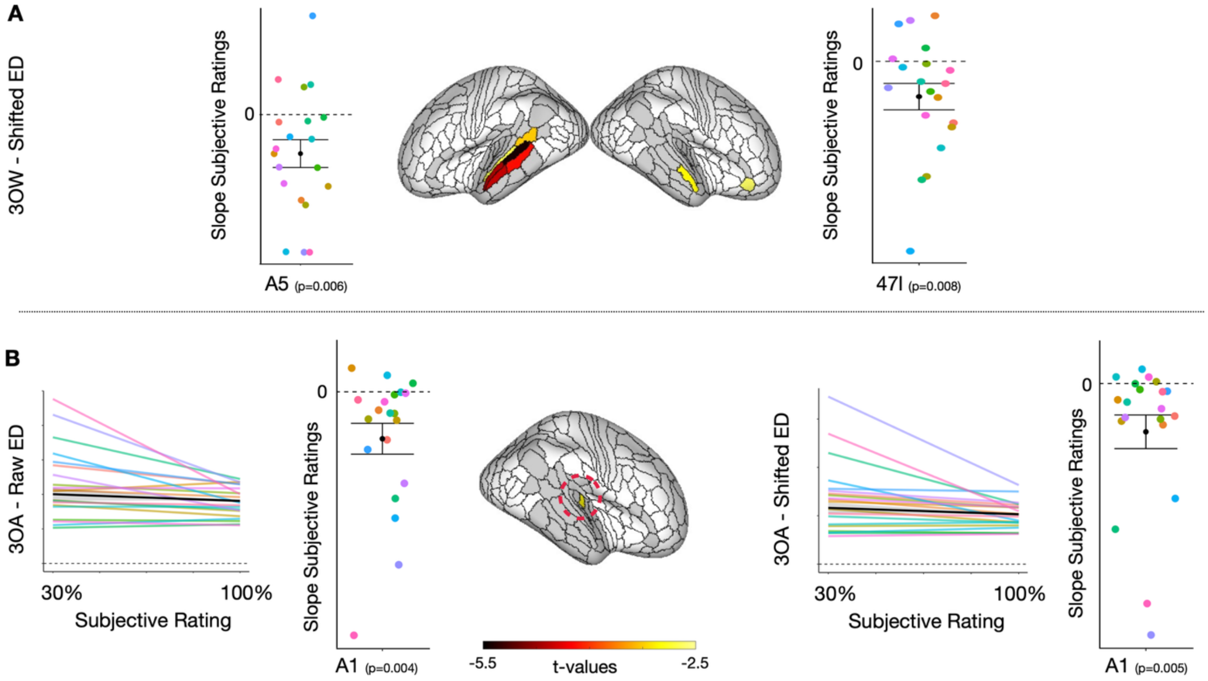


**Fig. S4. Relationship between subjective ratings and Euclidean Distance (ED), uncorrected results (p < 0.01). A. Uncorrected results for experiment 3OW after centroid alignment.** The spatial map shows the spatial distribution of significant uncorrected effects and the individual subject slopes. **B. Uncorrected results for experiment 3OA using raw data (left) and**

**data after centroid alignment (right).** For selected uncorrected significant regions, we display the regression lines and individual subject slopes. Gray-colored ROIs denote all brain regions included in the analysis.

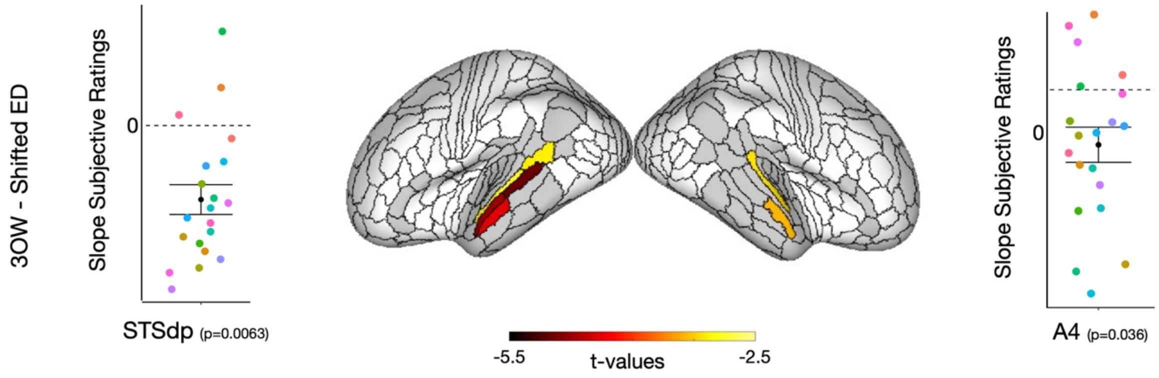


**Fig. S5. Subjective rating predicts the Euclidean Distance (ED), with a variance explained of at least 70% (FDR-corrected results).** Regions in which the subjective rating significantly predicts the ED, with a general trend of decreasing distance between the attended and the object alone representation, while increasing the performance (negative t-values). Here, we show the significant results for experiment 3OW, using the ED after centering the PC scores by subtracting the category-specific centroid, thereby aligning the score to the centroid of experiment OA.
